# Mechanistical and structural basis of Kv channel inhibition by 4-aminopyridine

**DOI:** 10.64898/2026.09.12.751153

**Authors:** Bernardo I Pinto-Anwandter, Richa Agrawal, Trayder Thomas, Eduardo Perozo, Benoît Roux, Francisco Bezanilla

## Abstract

Inhibition of Kv channels by 4-aminopyridine (4AP) improves motor function in multiple sclerosis by enhancing neuronal excitability. The mechanism of inhibition and the structural basis of 4AP binding to Kv channels remain unclear. Here, we determined the structure of the Shaker V369I-I372L-S376T (ILT) mutant bound to 4AP at 3.3Å, demonstrating that 4AP binds to the closed state of the channel. This structure is inconsistent with an open channel block mechanism. Electrophysiology experiments show that 4AP binds even when intracellular pore access is constitutively blocked, suggesting that 4AP enters the pore through membrane-facing fenestrations. MD simulations and mutational analysis agree with the proposed fenestration pathway and suggest that 4AP binds in its neutral form. These results support a mechanism where 4AP binds to a partially activated closed state that prevents complete activation of Kv channels, explaining its pharmacological activity.

## Introduction

Voltage-gated K^+^ (Kv) channels play a crucial role in regulating neuronal excitability by controlling membrane repolarization^1-3^. Due to their participation in a variety of physiological processes from muscle contraction and neuronal communication to immunological response, Kv channels are the targets of many pharmacological modulators, especially in demyelinating diseases^4,5^. Multiple sclerosis (MS) is a chronic inflammatory disease of the CNS characterized by demyelination and resultant axonal conduction block^6^. In demyelinated axons, the change in electrical properties of the membrane and redistribution of voltage gated Kv channels impairs action potential propagation^7^. The Kv inhibitor 4-aminopyridine (4AP), commercialized as dalfampridine, helps overcome conduction block ^8^. Dalfampridine is approved to improve walking speed and lower-limb strength in MS patients by enhancing neural conduction through demyelinated regions^9,10^. The clinical benefits of 4AP underscore the importance of Kv channels as therapeutic targets in demyelinating diseases. A detailed mechanistic and structural understanding of how this drug interacts with Kv channels would be highly desirable for rational drug design.

Current consensus suggests that 4AP inhibition takes place only when the channel activates. Thus, a sequential set of depolarizing pulses are required to observe the full effect of the drug, suggesting an open pore block mechanism^11,12^. Additionally, 4AP prevents the complete activation of the voltage sensor domain (VSD), reducing gating charges by about 10% ^13 14^. The effects on VSD movement are reminiscent of the ILT mutation that stabilizes the closed state of the channel with a partially activated VSD^15,16^. Therefore, once 4AP binds it both, blocks permeation and promotes channel closure^17^. In this open channel block and closed state stabilization model, most voltage-sensor movements representing the initial gating charge movement proceed normally, but the final channel-opening transition is impeded by 4AP stabilizing the closed state. However, alternative models have been proposed where 4AP preferentially binds to and stabilizes a closed partially activated or “intermediate” state of the channel, rather than acting as an open channel blocker^11^.

Here, we show that 4AP binds to the ILT mutant in the closed state when the VSD is partially activated. Moreover, the cryo-EM structure of the Shaker ILT mutant channel in complex with 4AP strongly suggests that it binds into the closed pore. Effects of 4AP on the S4 movement were observed even when access to the pore cavity was constitutively blocked, indicating that 4AP likely accesses the pore through membrane-facing fenestrations. Molecular dynamics (MD) simulation analysis revealed that changes in fenestrations at the interface of the pore helices allow 4AP unbinding. Changing the size of a residue at the inter subunit interface of the pore eliminates the use dependency of 4AP binding in the ILT mutant, supporting the existence of fenestrations in the closed state. The model presented here reconciles experimental observations providing a mechanistic and structural basis for 4AP inhibition of Kv channels.

## Results

### 4AP binds to the partially activated closed channel

To examine how 4AP interacts with distinct conformational states of the Shaker potassium channel, we compared its effects on the wild-type (WT) and the ILT mutant. In WT channels, when the channel was kept closed by holding at −80 mV for 30 minutes in the presence of 4AP, no inhibition was detected, indicating that 4AP cannot access its binding site while the channel remains in the closed state (**Fig. 1A**). However, after repeated depolarizations that open the channel the reduction in current amplitude caused by 4AP inhibition becomes apparent. The ILT mutation stabilizes a particular state of the closed channel in which three of the four voltage sensing arginines R1-R3 have translocated across the hydrophobic plug, but not R4 (R4 down)^16,18^. When 4AP was applied to ILT channels at 0 mV, a potential that maintains the channels predominantly in the R4 down closed state, the inhibitory effect was observed even without prior depolarizing pulses (**Fig. 1B**). This indicates that 4AP can bind to the closed channel when the VSD is partially activated (**Fig. S1**). Consistent with these results maintaining a -90 mV hyperpolarized voltage while applying 4AP in the ILT mutant prevents the closed state inhibition (**Fig. S2**). In the WT channel, 4AP reduces the total gating charge by approximately 10%, indicating that the VSD fails to completely activate once 4AP is bound. In the ILT mutant, about 10% of the gating charge is displaced to very positive potentials, reflecting the existence of a late voltage-dependent component associated with the final movement of the voltage sensor. When 4AP was applied to ILT, this component disappeared (**Fig. 1C**). These results indicate that 4AP stabilizes the closed state observed in the ILT mutant.

**Fig 1.**
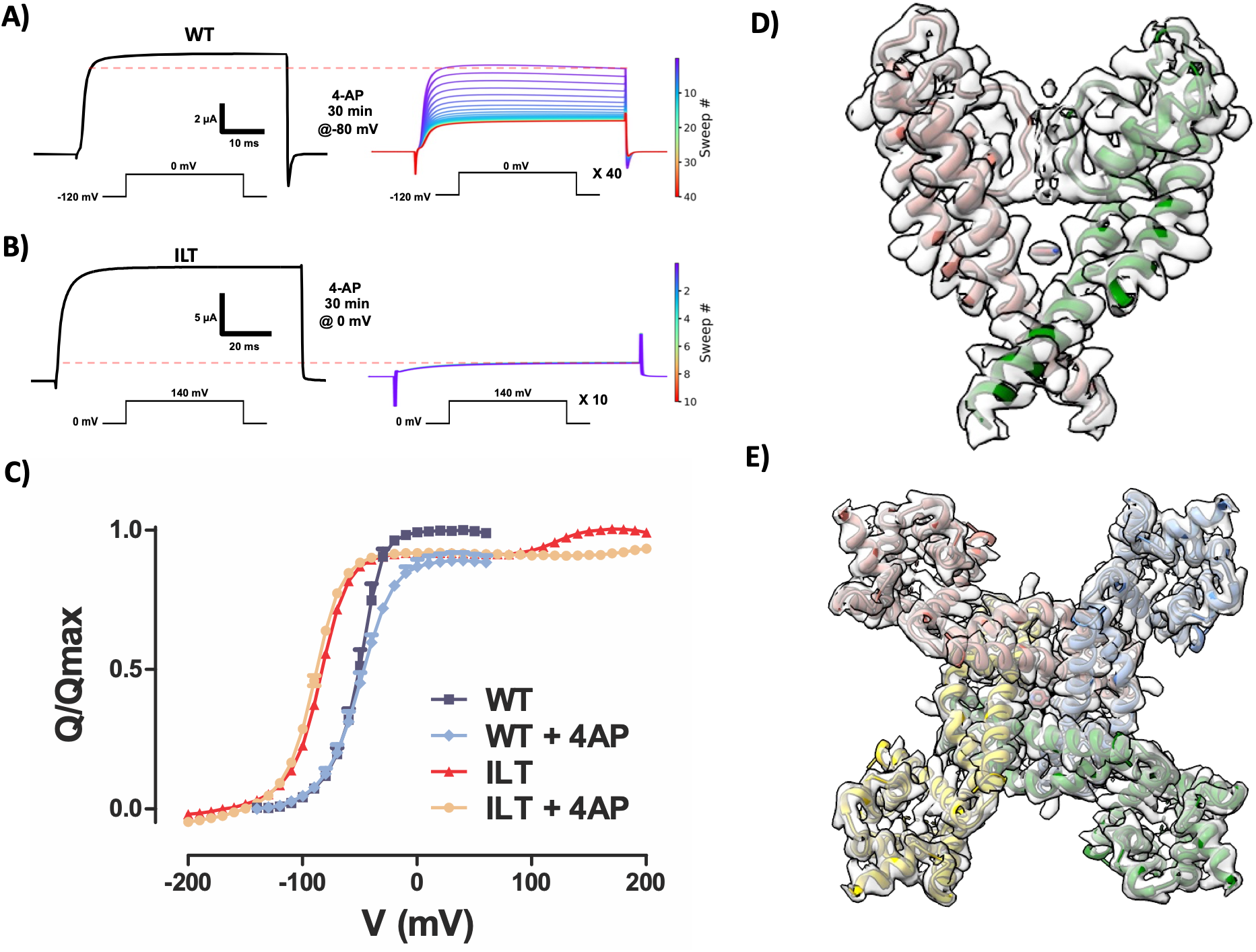
4AP binding in the Shaker ILT mutant. **A)** Shaker IR channel (WT) currents in response to a voltage pulse before (left) and after (right) 30-minute incubation with 400 µM 4AP while holding the membrane voltage at -80mV. Discontinuous red line indicates current level at the end of the first pulse after 4AP application, colors indicate the order of sweeps applied every 5 seconds. **B)** ILT mutant currents in response to a voltage pulse before (left) and after (right) 30 minutes incubation with 200 µM 4AP while holding the membrane voltage at 0 mV. **C)** Q-V curve of the WT (dark blue) and ILT mutant (red) and in the presence of 5 mM 4AP (sky blue WT, orange ILT). The gating currents were obtained by introducing the W434F background mutation. **D**) Shaker ILT-4AP model within unsharpened map showing pore helices of two chains colored in salmon, and green (membrane view); 4AP map density (gray) captured within the pore between S6 helices. **E**) Shaker ILT-4AP tetramer model colored in salmon, cornflower blue, green and gold within unsharpened map (intracellular view); 4AP is colored is brown color.

### Structure of the Shaker ILT mutant in the presence of 4AP

Based on our electrophysiological results, we attempted to determine the ILT structure in the presence of 4AP using single-particle cryo-EM and resolved the structure at 3.32 Å resolution (**Fig. S3**). We observed a clear map density corresponding to 4AP in the pore region of the tetrameric channel (**Fig. 1D, 1E, Fig. S4**). Overall, the structure of ILT–4AP shows key differences from the ILT structure (**Fig. 2A, B**). Notably, conformational changes were observed in the S4– S5 linker, as well as in the pore S5 and S6 helices. Similar to the ILT structure, the S4 helix in ILT–4AP remains in a partially activated state; however, the S4–S5 linker is shifted upward approximately ∼3.5 Å compared to the ILT structure (**Fig. 2C**). In the pore domain, the ILT–4AP structure shows approximately ∼1.8 Å shift in both the S5 and S6 helices away from the central axis of the pore at the intracellular side of the channel (**Fig. 2D)**. We compared the ILT–4AP structure with other previously reported Shaker channel structures, including the WT (PDB ID: 7SIP) (**Fig. S5A & B**) and the I384R mutant (PDB ID: 9OIC) (**Fig. S5C & D**), and found that ILT–4AP adopts a distinct conformation from those reported previously. The ILT–4AP structure shows an intermediate displacement of the S4-S5 linker compared to Shaker ILT and Shaker I384R (**Fig. S5E**). Similarly, the pore domain at the intracellular side shows a conformation that is in between ILT and I384R; this is also observed when plotting the pore radius (**Fig. S5F & G**). The selectivity filter region shows full occupancy of potassium ions with intact hydrogen bond interactions, indicating a conductive conformation as observed in the ILT structure (**Fig. S6**).

**Fig 2.**
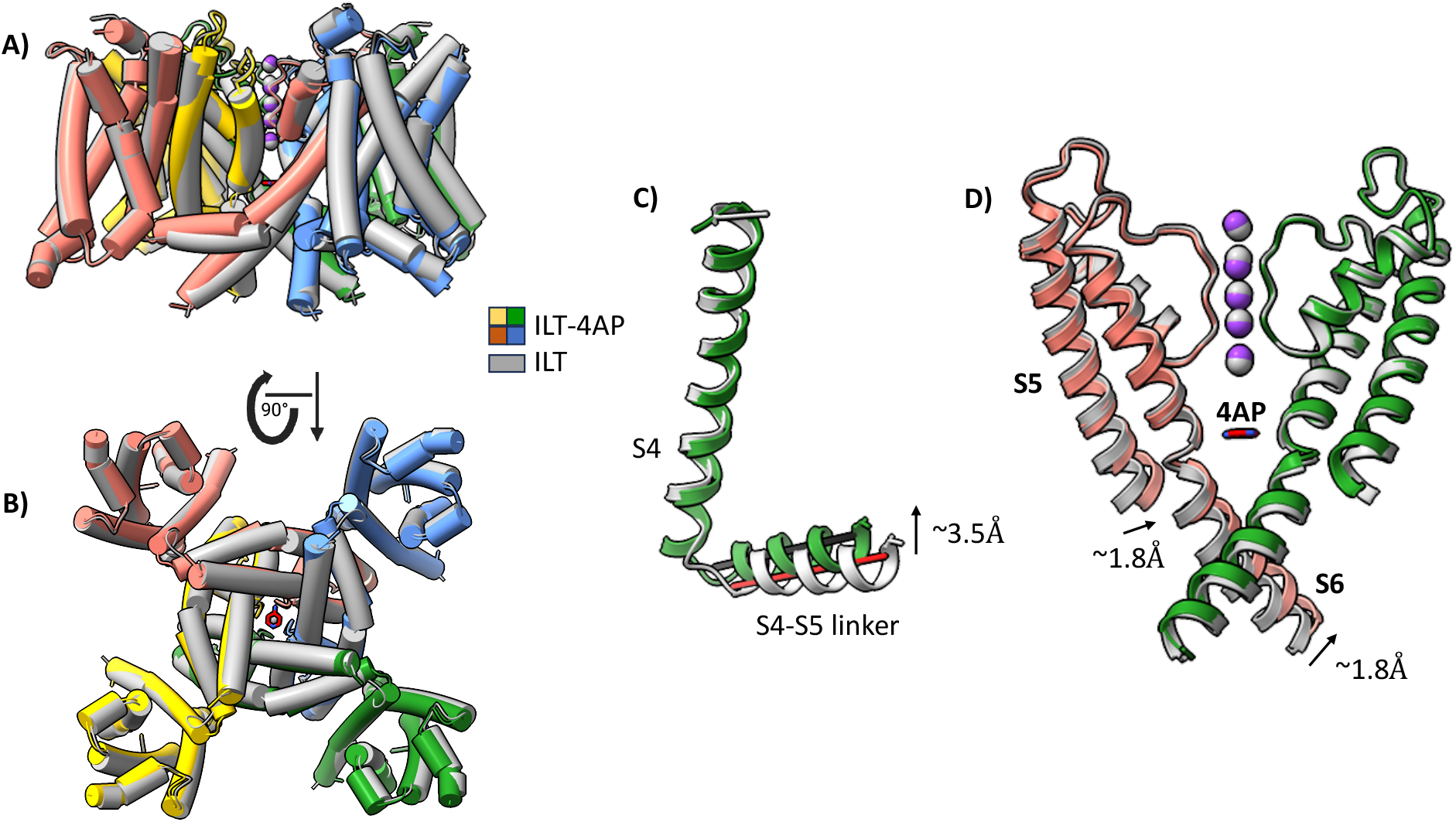
Conformational changes in Shaker ILT-4AP. A) & B) Membrane and intracellular view of Shaker ILT-4AP compared with Shaker ILT. Shaker ILT-4AP chains colored in salmon, cornflower blue, green and gold. Shaker ILT shown in gray. 4AP colored in brown and K+ ions are in purple C) Comparison of S4 and S4-S5 linker in Shaker ILT (gray) and Shaker ILT-4AP (green) showing that S4 position remains unchanged and S4-S5 linker is shifted. D) Comparison of S5 and S6 pore domain helices in 2 chains of Shaker ILT (gray) and Shaker ILT-4AP (salmon and green) where a small shift is observed.

### Binding site of 4AP within the pore

In the ILT–4AP structure, 4AP is observed bound within the pore region in a binding pocket formed by the pore S6 helices between residues I470 and V474 (**Fig. 3A**). This pocket is lined by I470, A471, and V474 where the ligand is stabilized through a combination of hydrogen bonding and hydrophobic interactions (**Fig. 3B, C**). Notably both the pyridine and amino nitrogen of 4-AP form hydrogen bonds to the I470-carbonyl of two diagonally opposite chains. I470, A471 and V474 sidechains form hydrophobic interactions with the 4AP ring. At physiological pH 4AP exists mainly in a protonated state; however, increasing the pH of the solution increases its potency indicating that the neutral form is responsible for inhibition ^19^. Since our cryo-EM structure does not allow us to unequivocally assign the protonation state of 4AP, we performed MD simulations of both the neutral and protonated states of the ligand bound to the closed channel. For both states we observed the same interactions as in the cryo-EM structure but the hydrogen bond between the pyridine-nitrogen and the I470-carbonyl was instead mediated by a water molecule that entered through the fenestration (**Fig. S7 A & B**). For the protonated ligand the same water molecule additionally mediated a hydrogen bond to the backbone carbonyl of V467. The dynamics of 4AP were observed to differ significantly between protonation states, the neutral ligand readily rotated in the membrane plane, while the protonated ligand was relatively conformationally stable and remained in the same quadrant (**Fig. S7 C & D**). To quantify the differences in binding between protonation states we performed alchemical free-energy calculations (**Fig. 3A, Fig. S7 & S8**) to estimate the binding affinity of each state to the closed conformation. The calculated binding energy for charged 4AP is +10.06 ± 5.43 kcal/mol (≥ +4.63 kcal/mol), indicating that it effectively does not bind. In contrast, the calculated binding free energy for neutral 4AP is -5.15 ± 0.06 kcal/mol, corresponding to a *K*_d_ of 176 µM. However, because the pKa of 4AP is 9.11, only a small (∼2%) population of 4AP is neutral at pH 7.4, therefore requiring a ∼50× higher concentration in solution to achieve 50% occupancy. Correcting for the protonation equilibrium, the binding free energy is -2.82 kcal/mol (Kd = 9210 μM). In line with the observed interactions, mutations of I470 to alanine or cysteine greatly diminish the effect of 4AP over the channel indicative of a decrease in binding (**Fig. 3D, E & F**).

**Fig 3.**
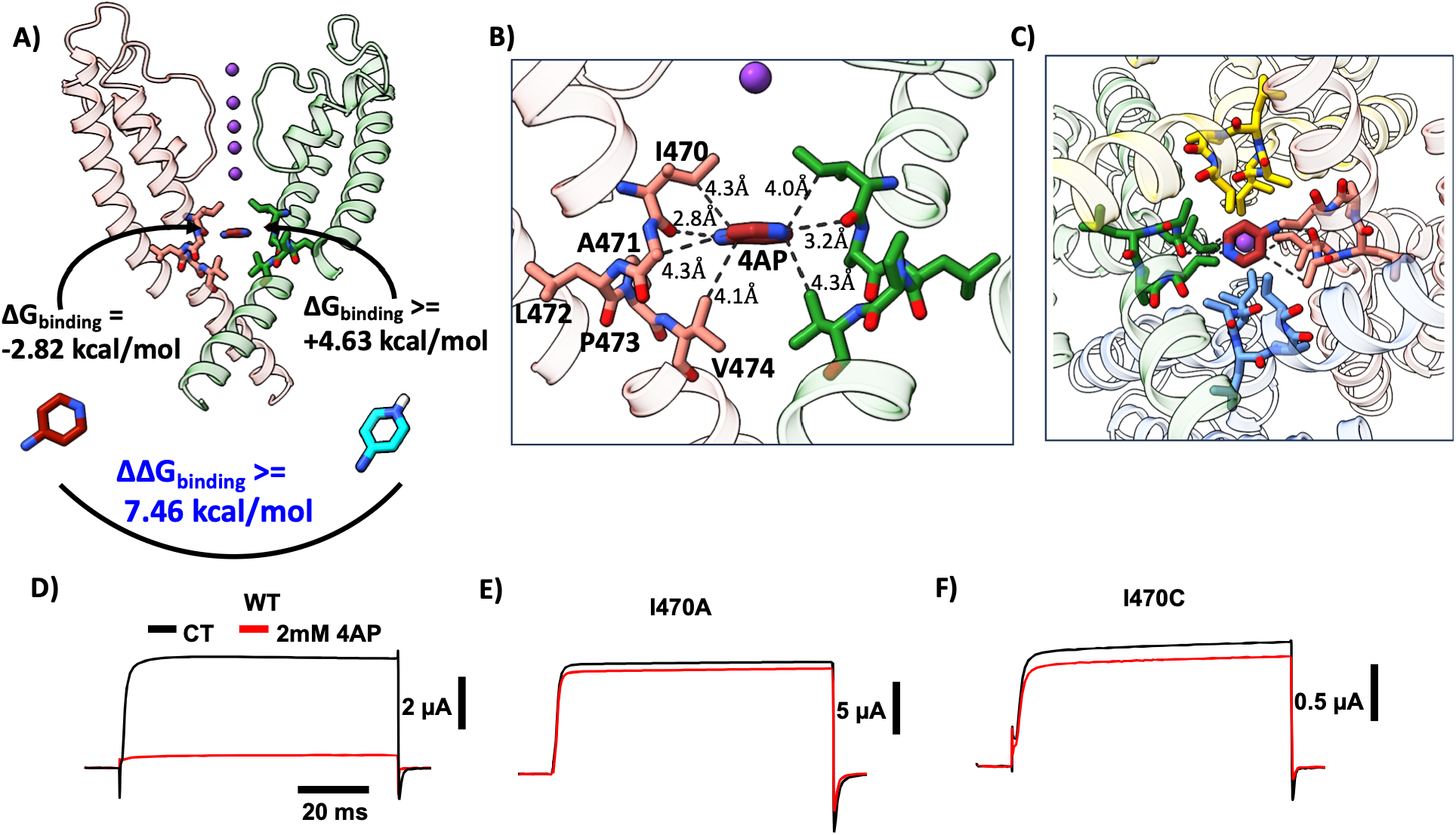
Interaction of 4AP in the pore. **A) Membrane** view of the Shaker ILT-4AP structure showing pore helices of two opposed chains (salmon & green); 4AP (brown) is shown at the pore region between S6 helices. 4AP binding pore lining residues (I470-V474) are shown as sticks. **B)** Zoomed membrane view of pore lining residues I470, A471 and V474 of S6 helices showing interactions with 4AP. **C)** Zoomed intracellular view of pore lining residues I470, A471 and V474 of S6 helices showing orientation of 4AP within pore. **D-F)** Currents in response to a voltage pulse before (black) and after (red) application of 2mM 4AP for the (**D**) WT, (**E**) I470A and (**F**) I470C mutants.

### Binding of 4AP does not occur through the permeation pathway

Despite observing binding in the permeation pathway, it is not completely clear from the structure how 4AP enters its binding site. It is possible that rearrangements among closed states of the activation gate analogous to what we observe in ILT-4AP might allow 4AP to enter from the intracellular side into the binding site. Alternatively, since 4AP can partition into the membrane it is possible that it can diffuse along the membrane into the binding site through a fenestration between the S6 helices. To test if 4AP requires binding through the permeation pathway we used the V478W and W434F mutations (**Fig. 4A, Fig. S9)**. The V478W mutant produces a non-conductive channel by replacing a small hydrophobic residue along the conduction pathway with a bulky tryptophan preventing intracellular access to the internal cavity and eliminating potassium permeation^20,21^. Since the V478W channel is non-conductive we can observe the effects on the gating currents and compare them to the gating currents of the W434F channel. The W434F mutant produces a non-conductive channel by enhancing the slow inactivation process through changes in the selectivity filter without changes in the internal cavity accessibility^22-24^. If 4AP diffuses from the intracellular side, the V478W mutation would prevent its binding. Contrary to this, in both mutants 4AP produces the same 10% decrease in total gating charge (**Fig. 4B**). The addition of 4AP also produces an acceleration of the gating currents deactivation (**Fig. 4C & D**). This acceleration occurs because VSD deactivation is slower in open channels than in closed channels and since 4AP prevents opening the deactivation of the gating currents is accelerated, producing larger currents despite having less total charge^24,25^. These results show that 4AP binding does not occur from the intracellular side but is likely through fenestration between S6 helices.

**Fig 4.**
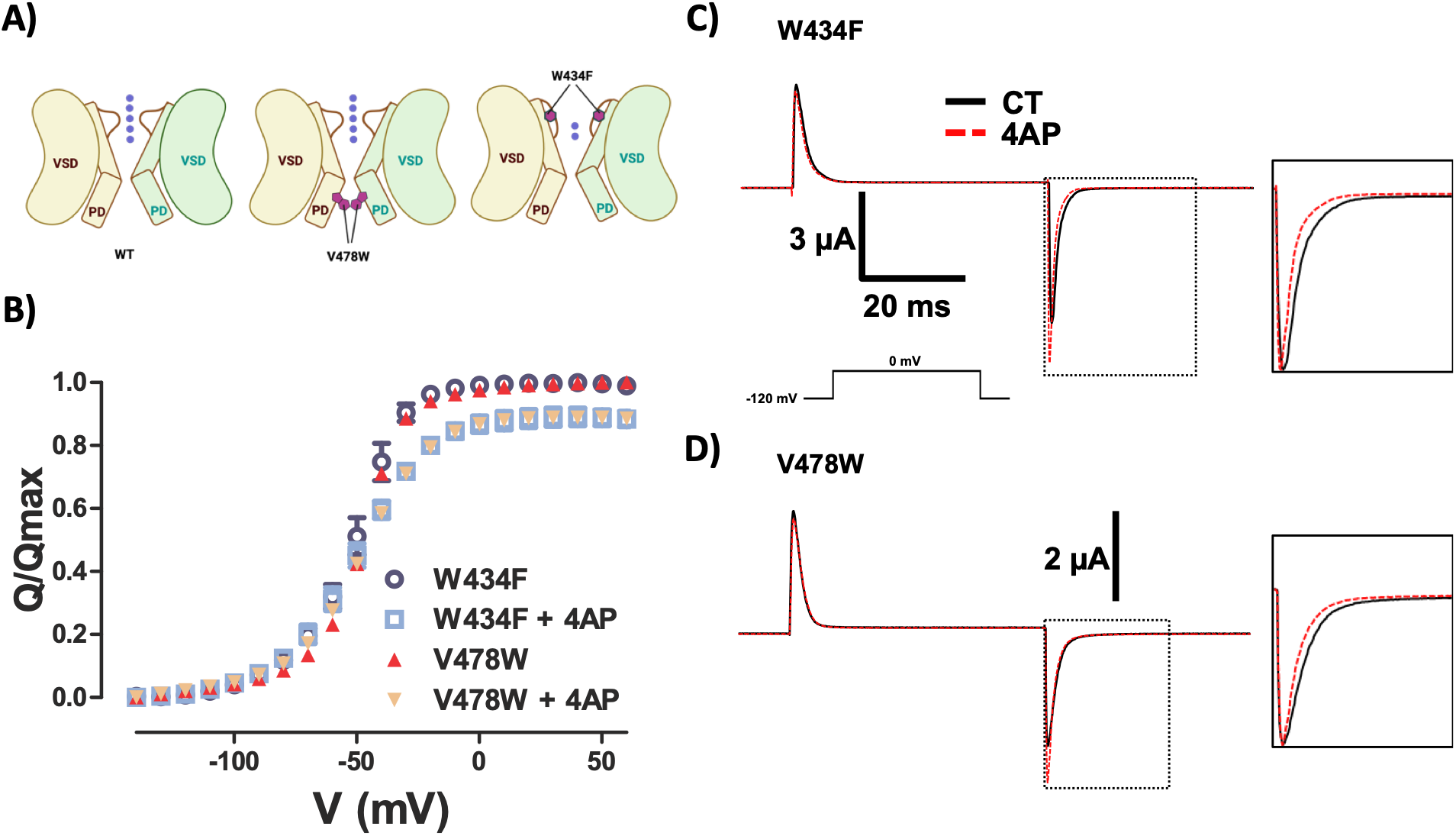
Binding of 4AP to permeation blocked channel. **A)** Diagram of the permeation pathway in the WT and illustration of the effects of the V478W and W434F mutations. **B)** Q-V curve of W434F (open symbols red) and V478 (filled symbols, blue), and in the presence of 5 mM 4AP (W434F open symbols orange, V478W filled symbols sky blue). **C, D**) Gating currents in response to a voltage pulse before (black continuous line) and after (red discontinuous line) application of 5 mM 4AP for W434F (**C**) and V478W (**D**) mutants. The squares show normalized deactivation currents of the selected region to illustrate the deactivation acceleration.

### Binding of 4AP through membrane facing fenestrations

As observed in the ILT structure (PDB ID: 9ZS7), entry of 4AP from the intracellular side is restricted; however, a small access pathway is present between the S6 helices. An alternative route into the pore may occur through membrane-facing fenestrations formed by residues V467, V468, and P473 of the S6 pore helices (**Fig. 5A & B**). Notably, the conformational changes observed in the ILT–4AP structure are localized near this entry point and involve repositioning of residues L396 and I400 in the S5 helix. These rearrangements likely contribute to trapping 4AP within the central cavity, preventing its exit. We performed molecular dynamics simulation of the ILT-4AP structure and focused on this possible fenestration pathway. By performing steered MD simulations, we observed a 4AP unbinding event where 4AP leaves the binding site through the proposed fenestration. This unbinding event occurs alongside a temporary helical shift around P473 in the S6 kink region (**PXP motif**) (**Fig. S10**). These results provide evidence for a fenestration mediated entry of 4AP into the pore cavity. Among the fenestration residues identified, previous studies have shown that increasing V467 side-chain size (V467I) significantly reduces the apparent affinity for 4AP, despite it not being part of the binding pocket ^18^. This indicates that V467 plays a key role in controlling access of 4AP to its binding site. To study the role of V467 in allowing 4AP access to its binding site we focused on the use dependence in WT and ILT. Similar to the WT, the ILT mutant exhibits a clear dependence on pulse cycle period, (**Fig. 5C & D, Fig. S11**). Taken together, these observations suggest that accessibility to the binding site increases during channel activation by increasing the fenestration entry point. Mutation of V467A in ILT (ILT–V467A) abolishes the dependence of 4AP binding kinetics on cycle period from the partially activated state (**Fig. 5E, Fig. S12**), without significant changes in voltage dependence (**Fig. S13**). Thus, by reducing the size of V467 we increase the size of the fenestration in a way that the binding site becomes equally accessible in the partially activated closed state than during activation. Taken together these results support our fenestration accessibility hypothesis.

**Fig 5.**
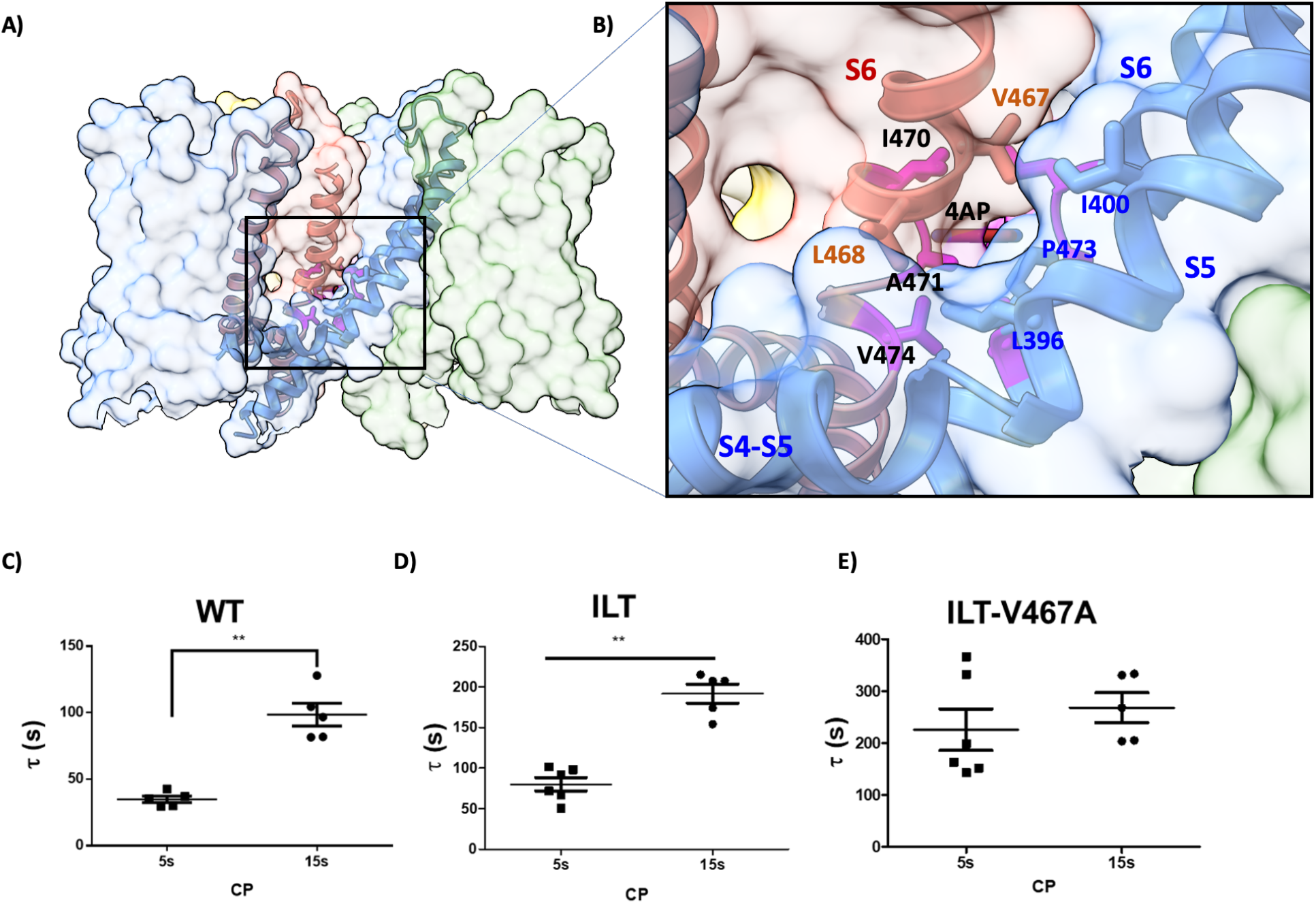
Fenestration access of 4AP into the pore. A) Surface view of Shaker ILT-4AP tetramer where S4-S5 linker and pore helix of two chains are shown in the ribbon form colored in salmon and cornflower blue. Fenestration entry point formed by S6 helix residues (V467, A471, P473, L468) and covered by S5 helix residues (L396 and I400) is shown. B) Zoomed view of all S6 helix residues (V467, A471, P473, L468) and S5 helix residues (L396 and I400) which form the fenestration entry point, residues which is part of binding highlighted in magenta color, 4AP in brown color **(C-E)** Effect of cycle period on the inhibition time constant of 4AP for WT (400 µM) ILT (200 µM) and ILT-V467A (1 mM) mutants. ** p<0.01 Mann-Whitney U test.

## Discussion

### A structural model for 4AP inhibition

Based on the present results, we propose an explicit mechanistic model for 4AP inhibition of Kv channels (**Fig 6A**). The model includes two closed states and one open state, where the first transition among closed states state (from R1 down to R4 down) represents the translocation of gating charges R1-R3 carrying 90% of the charge. Transition from the R4 down to open (R4 up) represents the translocation of R4 and carries a small fraction of the charge (10%). 4AP can bind into the R4 down state, stabilizing a conformation from which the channel cannot transit into the open state. In contrast with previously proposed models^17^, 4AP does not bind to the open state of the channel; as illustrated by the WT structure, in spite of the wider intracellular entry of the open channel (**Fig S14)**. 4AP binding site is only accessible in the R4 down state, and while 4AP does not bind to the R1 down state, the VSD can deactivate completely in presence of 4AP as indicated by the transitions between the R4 down and R1 down 4AP bound states.

**Figure 6.**
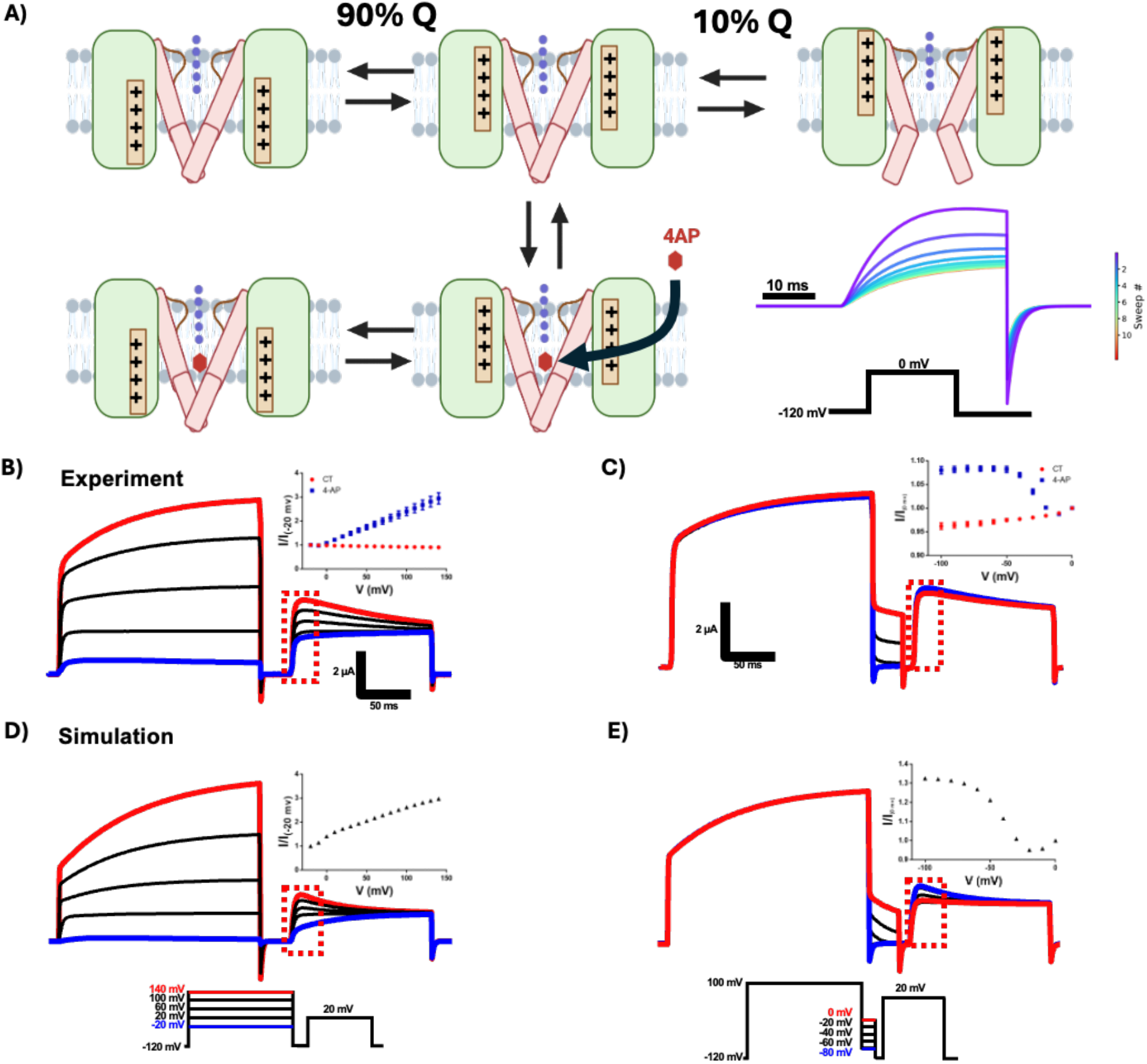
Structurally informed mechanistic model for 4AP action. **A)** Diagram of the 5-state model. Transition of the VSD from the resting closed sate (R1down) to the partially activated closed state (R4down) carry 90% of the charge movement, transition from R4down to the activated open state carries 10%. 4-AP binds only in the R4down state and after it binds can be trapped in the R1down state. The model reproduces the effect use dependence of 4-AP inhibition (lower right inset). **B**) Effect of repetitive activation after 4AP application (400 µM) on the simulated model ionic currents. **C**) Effect of depolarization on the unbinding of 4AP, inset graph shows the normalized peak current during the 20 mV step (square box) vs depolarizing voltage in the absence (red) and presence (blue) of 400 µM 4AP. **D**) Effect of variable repolarization after a large depolarization on the rebinding of 4AP, inset graph shows the normalized peak current during the 20 mV step (square box) vs repolarizing voltage in the absence (red) and presence (blue) of 400 µM 4AP. **E**) and **F**) are simulations of the experiments in **C** and **D** respectively using the 5-state model with 400 µM 4AP.

This model reproduces the observed use dependence of 4AP inhibition (**Fig 6A, inset**). In the model, since going from the partially activated closed state to the open state is voltage dependent, a large membrane depolarization would drive channels into the open state, causing 4AP to unbind. Applying large depolarizing pulses, leads to a clear second slow kinetic component associated with 4AP unbinding (**Fig 6B**). Subsequently, if after a depolarization that unbinds 4AP we apply a strong hyperpolarization, the channels would rapidly move to the resting closed state, passing only briefly through the partially activated state. This short visit does not allow enough time for 4AP to rebinding events. Consequently, if we apply a second depolarizing pulses, the channels that were relieved from inhibition (red trace in **Fig 6B**) will show a faster activation and larger currents followed by a reestablishment of 4AP inhibition (**Fig 6B**). Additionally, a strong depolarization followed by different levels of hyperpolarization produces progressively larger test currents with increasing hyperpolarization, consistent with reduced rebinding at more negative voltages (**Fig 6D**). We can reproduce these effects using our simplified gating model (**Fig 6E, F**). The similarity of the observed effects in the channel and the simulations from this minimal five state model is outstanding, providing a clear structurally informed mechanistic basis for 4AP inhibition.

### Insights into 4AP binding and dynamics

Free-energy calculations favor a mechanism where 4AP binds in its neutral form. This observation can be largely explained by the considerable solvation free energy of the charged form in solution, -52.1 kcal/mol compared to -8.0 kcal/mol for the neutral form. Binding of the neutral form is consistent with the observation that the apparent affinity of 4AP improves at more negative voltage^26^. This is opposite to what would be expected for a positively charged pore blocker and may also have implications when considering a membrane fenestration binding model. Though difficult to measure experimentally, an effective *K*_d_ for the closed state can be estimated from the observation of McCormack et al. where 100 µM of 4AP produced a 95% inhibition of the WT current at -40 mV, a voltage at which the channels predominantly occupy the closed state ^26^. Assuming that the extent of inhibition reflects equilibrium binding, this corresponds to an effective *K*_d_ (closed) of approximately 5.26 µM. However, this calculated binding affinity is weaker than this experimental estimate perhaps due to inaccuracies in the model. Evaluation of the interaction energy of 4AP in the binding site using an *ab initio* QM approach indicates that the current force field model underestimates the interaction energy by ∼1 kcal/mol. But this does not considers the slow kinetics of the underlying processes.

### The nature of the partially activated and resting closed states

Our work explains several well-known physiological effects of 4AP on Kv channels, including its use dependence, the decrease in total gating charge, and the acceleration of deactivation gating currents. More broadly, our findings provide insight into the nature of the closed states that precede opening. Our results predict that the S6-fenestration in the pore changes between closed states in a way that governs 4AP access. The use dependence and persistence of inhibition after washout indicates that 4AP does not associate or dissociate freely in the resting state of the channel^12^. This is consistent trapping of 4AP in the resting state by closure of the fenestration, thus relieve from inhibition requires voltage-dependent transitions that reopen the fenestration exit pathway allowing 4AP to unbind. This means that different closed states are not distinguished solely by changes in the VSD but also by structural rearrangements in the pore. In a similar fashion, accessibility to cysteine modifying agents of the V478C mutant located in the intracellular mouth of the channel is increased in the partially activated VSD compared to the resting state, indicating structural changes in the pore between closed states^27^. Overall, this evidence indicates that VSD transitions along closed states produce conformational changes in the pore.

### The role of fenestrations in Kv channels

Fenestrations are a common structural feature through which lipids and small molecules interact with the pore of ion channels, usually after partitioning into the low dielectric regions of the lipid bilayer. In voltage-gated sodium channels, fenestrations provide a lateral pathway from the membrane to the central cavity and strongly influence resting-state block. In NavAb, changing the size of the fenestration by mutations produced changes in the potency of local anesthetics inhibition^28^. Recent cryo-EM structures of human Nav1.7 have revealed differential binding of several antagonists through fenestrations, with vinpocetine and hardwickiic acid occupying the repeat III–IV fenestration and vixotrigine interacting with the repeat IV–I fenestration^29^. Cryo-EM structures of CaV channels show that fenestrations participate in ligand binding. In CaV3.1, the selective antagonist Z944 occupies the central cavity by transiting through the repeat II–III fenestration; in CaV3.2, ACT-709478 and TTA-A2 extend into the repeat IV–I fenestration, whereas TTA-P2 and ML218 extend into the II–III fenestration; and in CaV1.1, the dihydropyridines nifedipine and Bay K 8644 bind in the III–IV fenestration^30-32^. In the Ca^+2^ activated K^+^ channel BK membrane-facing fenestrations are also observed in Ca^+2^ free BK structures. Work on the prokaryote homolog MthK channel indicates that these fenestrations allow the accessibility of quaternary ammonium derivatives to the internal cavity in the closed pore ^33^, arguing for an ancient origin to this inhibitory mechanism. More broadly, fenestration-associated ligand binding has also been described in K2P ^34^ and KCNQ ^35^ channels, supporting the idea that membrane-facing fenestrations can mediate drug access and lipid interactions across multiple ion-channel families.

## Material and methods

### Cell lines

The HEK293S GnTI-cells in suspension that were used for protein expression and purification were obtained from ATCC (CRL-3022). GnTI-cells were grown at 37 °C and 7.8% CO_2_ in FreeStyle 293 expression medium (Gibco, Thermo Fisher Scientific) supplemented with 2% heat-inactivated fetal bovine serum (FBS) and 10 µg ml^−1^ penicillin–streptomycin. Sf9 cells (Thermo Fisher Scientific, 12659017) were cultured in SF-900 II SFM medium (Gibco, Thermo Fisher Scientific) supplemented with 10% FBS and 10 µg ml−1 gentamicin at 28 °C.

### Cryo-EM sample preparation and data acquisition

Shaker ILT plasmid construct, and protocol used for expression and purification has been described previously ^16^. Freshly prepared concentrated protein ∼1 mg ml^−1^ was used for cryo-EM. To solve structure with 4AP (4-Aminopyridine), 10mM concentration (150mM stock) is used in respective buffer. UltrAufoil 300-mesh 1.2/1.3 Au grids (Quantifoil) were plasma-cleaned at 40W for 30 s in an air mixture in a Solarus Plasma Cleaner (Gatan). Purified Shaker ILT samples mixed with 10mM 4AP and incubated for 1 hour on ice, were applied to the grids and frozen in liquid-nitrogen-cooled liquid ethane using a Vitrobot Mark IV (FEI) and the following parameters: 3.5 µl sample volume; 3.5 s blot time; 1.0 blot force; 100% humidity; temperature of RT. Grids were screened for single particles on a 200kV Glacios TEM with cryo-autoloader equipped with a K2 summit direct detector (Gatan). Grids were imaged at the University of Chicago on a Titan Krios with a K3 detector and GIF energy filter (set to 20 eV) at a nominal magnification of ×81,000, corresponding to physical pixel size of 1.068 Å. Movies were acquired for 50 frames with total exposure 60 e^−^ A^−2^.

### Single-particle cryo-EM analysis

All steps for structure determination were performed using CryoSPARC ^36^, including motion correction and contrast transfer function (CTF) estimation. A dataset of ∼10,700 movies was collected, and particles were picked and classified in 2D to generate templates for template-based particle picking. Approximately ∼2 million initial particles were picked and subjected to 2 rounds of 2D classification. From these, 0.35 million particles were selected to generate three *ab initio* models for C1 and C4 symmetry separately. This process was repeated multiple times to eliminate junk particles. Particles from the best class (∼0.25) were then processed for 3D refinement (non-uniform or local refinement) with C4 refinement algorithm^37^, which consistently yields best results. Reference-Based Motion correction was applied to estimate per-particle movement trajectories and empirical dose weights. Local resolution was calculated using CryoSPARC^36^.

### Model building, refinement, and molecular visualization

The Shaker-IR ILT structure (PDB ID: 9ZS7) was used as a template to build atomic model of Shaker-ILT-4AP. We used phenix_eLBOW program in Phenix for generation of 4AP ligand restraints and optimization^38^. Secondary structural elements were fit into the map and further model building was pursued through iterative rounds of manual model building in COOT ^39^ registering secondary structural elements using bulky residues such as Phe and Arg. Simultaneously, flexible loops fitting in position in the atomic models were obtained using interactive flexible fitting in ISOLDE^40^. The density was of sufficient quality to assign rotamers for key residues whereas side chains of some residues that could not be assigned even tentative rotamers were truncated at the Cβ position of the residue. The tetramer model was generated by applying C4 symmetry operations to the monomer in UCSF ChimeraX^41,42^. Models were refined iteratively using phenix.real_space_refine in Phenix^43^. PDB validation has been performed in the PDB validation server. All structural analyses and figure generation were performed using UCSF ChimeraX.

### Site-directed mutagenesis and electrophysiological recordings

*Xenopus laevis* ovaries were obtained from Xenopus 1 (Dexter, Michigan). The follicular membrane was digested by collagenase 2 mg/mL supplemented with bovine serum albumin 1mg/mL. Oocytes were kept at 12 or 18 ºC in SOS solution containing (in mM) 96 NaCl, 2 KCl, 1 MgCl_2_, 1.8 CaCl_2_, 10 HEPES, pH 7.4 (NaOH) supplemented with gentamicin (50 mg ml^-1^). We used clones from Shaker zH4 K^+^ channel with removed N-type inactivation (IR, Δ6-46) in the pBSTA vector ^44^. Mutations were performed using Quikchange site directed mutagenesis and cRNA was transcribed from linearized cDNA, using T7 RNA kit. cRNA was injected in defolliculated oocytes (stage V-VI) and incubated in SOS solution at 18 or 12 ºC. After 1-4 days currents were recorded using the cut-open voltage-clamp method ^45^. Voltage-sensing pipettes were pulled using a horizontal puller (P-87 Model, Sutter Instruments, Novato, CA), and the resistance ranged between 0.2-0.5 MΩ. Data were filtered online at 20–50 kHz using a built-in low-pass four-pole Bessel filter in the voltage clamp amplifier (CA-1B, Dagan Corporation, Minneapolis, MN, USA) sampled at 1 MHz, digitized at 16-bits and digitally filtered at Nyquist frequency (USB-1604; Measurement Computing, Norton, MA) using Gpatch64M (in-house software). An in-house software (Analysis) was used to acquire and analyze the data. External solution for ionic recording was composed by (mM): K-Methanosulfonate (MES) 12, N-Methyl D-glucamine (NMG)-MES 108, Ca-MES 2, HEPES 10, pH 7.4, and internal solution by (mM): K-MES 120, EGTA 2mM, HEPES 10, pH7.4. In the case of mutants in the ILT background the external solution was adjusted to 120 K-MES and no NMG. External solution for gating currents recording was composed by (mM): NMG-MES 120, Ba-MES 2, HEPES 10, pH 7.4, and internal solution by (mM): NMG-MES 120, EGTA 2mM, HEPES 10, pH7.4. 4AP was diluted with external solution from a 200 mM stock to obtain the adequate concentration.

### Kinetic modeling

Simulations for the five-state model were carried out using IonChannelLab ^46^. From here we obtained the probabilities of each state with time for a given voltage and 4AP protocol. The modeling parameters and the scheme used are provided in **supplementary figure S15**.

### Molecular dynamics simulations

Molecular dynamics simulations were performed with NAMD 3.0.2, using the CHARMM forcefield, and CGenFF 4.6 to parameterize 4AP ^47-50^. To perform MD of the Shaker-ILT/4AP complex, a system was constructed from the cryo-EM structure (residue 206 to 491). Missing loops were modelled using AlphaFold 2 through ColabDesign v1.2.0b3 ^51,52^. Then the channel was embedded in a POPC bilayer with 150mM KCl and equilibrated according to the protocol of the CHARMM-GUI membrane builder ^53^. The ligand 4AP was added from the coordinates of the cryo-EM structure and the selectivity filter was loaded with potassium ions in sites S0,S2,S4 and water in sites S1,S3. An additional 100 ns NPT equilibration was performed with only the protein backbone restrained. Unbiased simulations were performed for 200 ns for both protonation states of 4AP bound to the ILT and ILT-V467A structures. Steered MD was performed on the neutral ligand from the equilibrated complex using the Colvars module ^54^. The distance was increased in the XY-plane between the center of masses of the 6 atoms of the 4AP ring and the CA atoms of residues 393-488 for a pair of adjacent protein subunits. The restraint distance was increased by 20 Å over 4 ns with a 10 kcal/mol force constant. Binding free energy calculations (Kd) for 4AP binding to the closed Shaker-ILT channel were conducted using the alchemical workflow of BFEE 3.2.1 ^55^, initiated from the equilibrated structure for both neutral and protonated forms of 4AP. The ligand and protein were treated flexibly, backwards, and forwards transformations were performed over 21 windows of 1 ns. In all bound simulations the ligand rotation in the xy-plane was restricted to a single quadrant of the binding site by additionally including a flat bottom restraint with a force constant of 10 kcal/degrees to the 90-degree wedge centered on the equilibrated pose. The ligand has 2-fold symmetry along its long axis, but only one of the two equivalent poses was sampled during simulations. The quadrant and symmetry restrictions were accounted for by a -*k*_B_*T*ln(4×2) adjustment to the calculated energy and an additional correction of −*k*_B_*T*ln(1⁄(1 + 10^p*Ka*−pH^)) (+2.33 kcal/mol) was introduced to account for the neutral ligand being only ∼2% of the total population, based on a pH of 7.4 and pKa of 9.11. To estimate error due to the force field, the interaction energy of 4AP to a cluster model of the binding site was evaluated in both QM and MM. The cluster model contained atoms within 4 of the ligands capped aliphatically, I470-CG1+CG2 to A471-CA including connecting backbone, and V474-CG2+CB. Protein coordinates were frozen, and the ligand was minimized separately in each of QM and MM. QM calculations were performed in Gaussian 16 Revision C.02 [https://gaussian.com/citation/] using the M06/def2-TZVP level of theory.

## ACKNOWLEDGEMENT

The authors are grateful to the staff of the single particle CryoEM facility at The University of Chicago for their assistance during sample screening and data collection. This work was supported by NIH grants R01GM030376 (FB), 5R35GM152124 (BR), 5R01GM150272 (EP) and by computational resources provided by the Research Computing Center of The University of Chicago and the Beagle3 high-performance GPU cluster funded by the NIH through grant 1S10OD028655. BP-A is a PEW Latin American Fellow.

## Author information

Author notes

These authors contributed equally: Bernardo I Pinto-Anwandter, Richa Agrawal

