## Supplementary Information for "Mechanistical and structural basis of Kv channel inhibition by 4-aminopyridine"

### Mechanistical and structural basis of Kv channel inhibition by the conduction enhancing drug 4-aminopyridine.

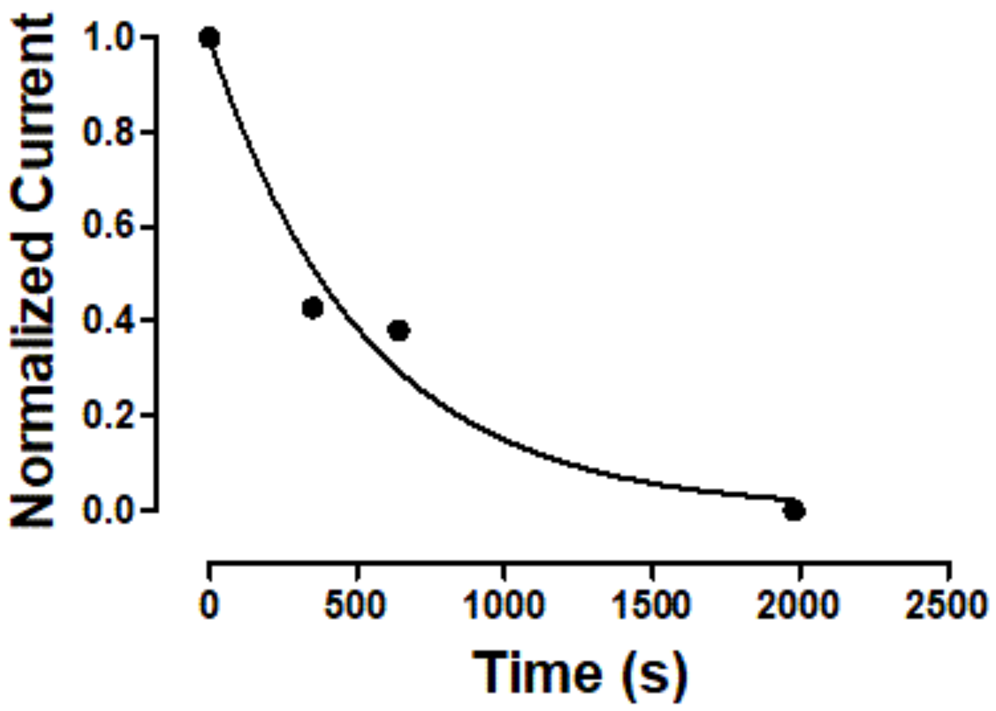


**Fig: S1. Time course of closed state 4AP binding to the ILT mutant**. Normalized current for the first pulse after incubation with 200 µM 4AP while holding the membrane voltage at at 0 mV. Currents were normalized by dividing by the current before 4AP application minus the baseline after steady state. The line shows a fitting to an exponential with time constant of ~560 s.

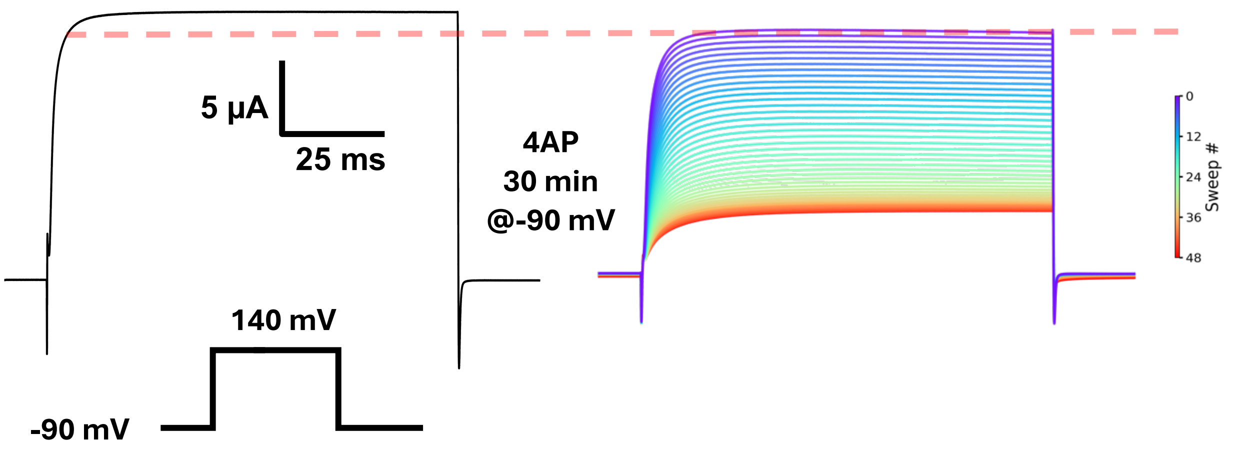


**Fig: S2. 4AP binding in the shaker ILT mutant in the resting state**. ILT mutant currents in response to a voltage pulse before (left) and after 30 minutes incubation with 200 µM 4AP while holding the membrane voltage at -90 mV. Discontinuous red line indicates current level at the end of the first pulse after 4AP application, colors indicate the order of sweeps applied every 5 seconds.

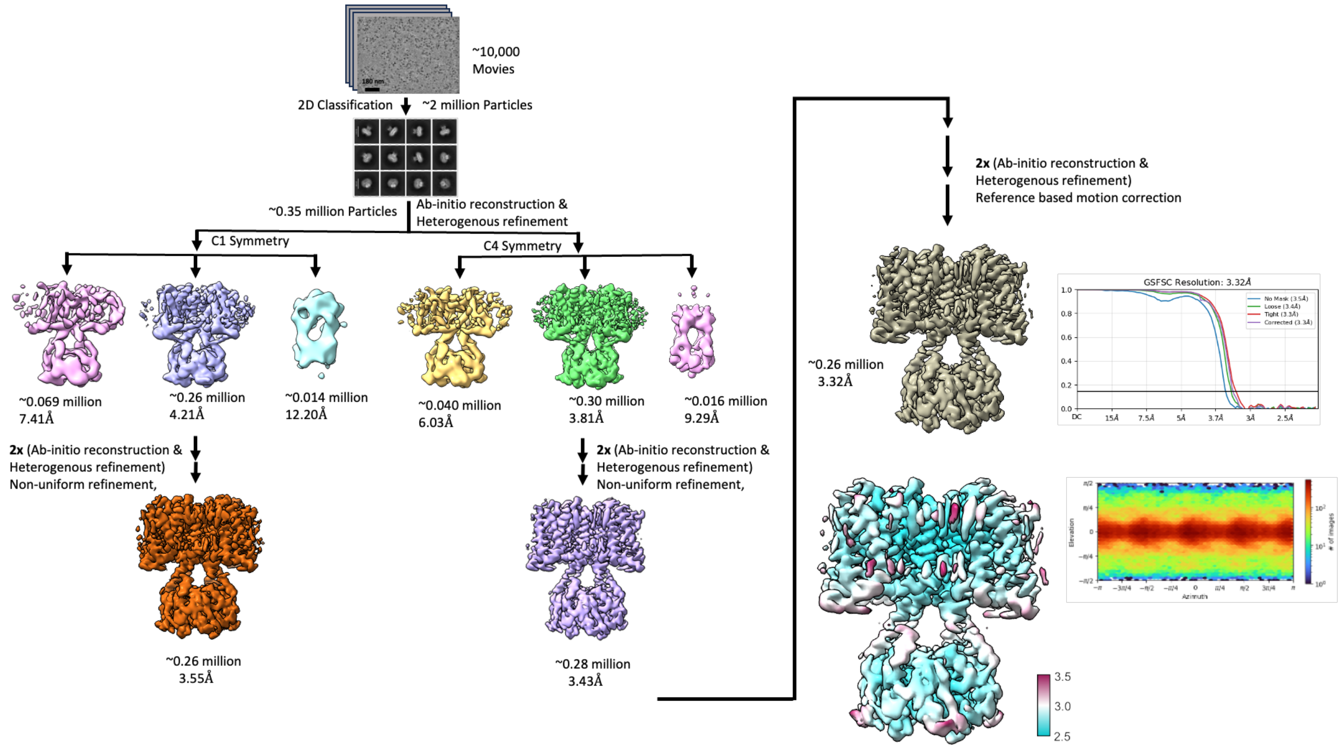


**Fig: S3. Data processing of shaker ILT-4AP**

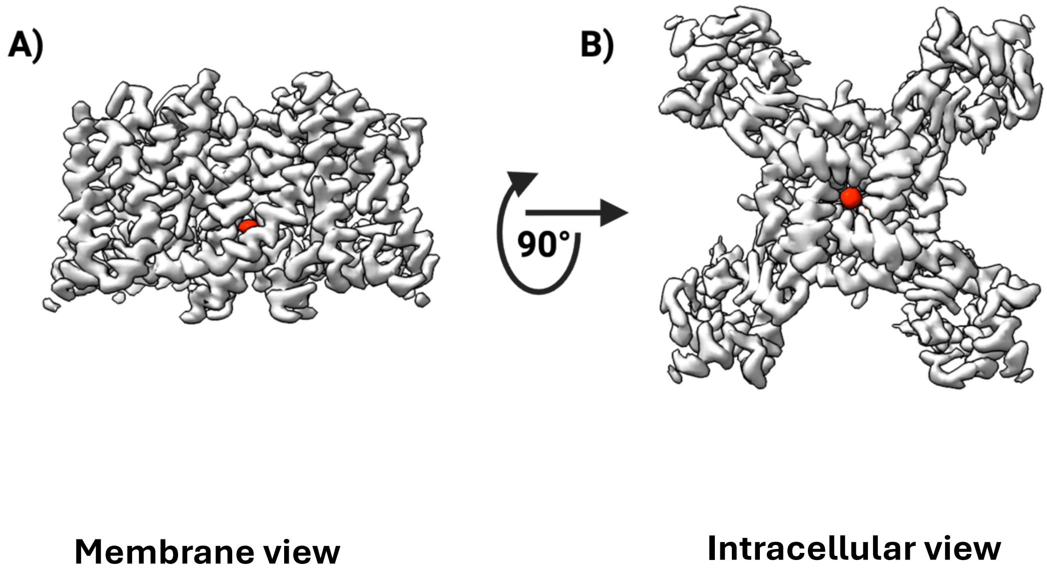


**Fig: S4. Tetramer map of ILT-4AP showing 4AP density**. A) Membrane view of tetramer unsharpened map shown in gray, 4AP map density in red. B) Intracellular view of tetramer unsharpened map shown in gray, 4AP map density in red.

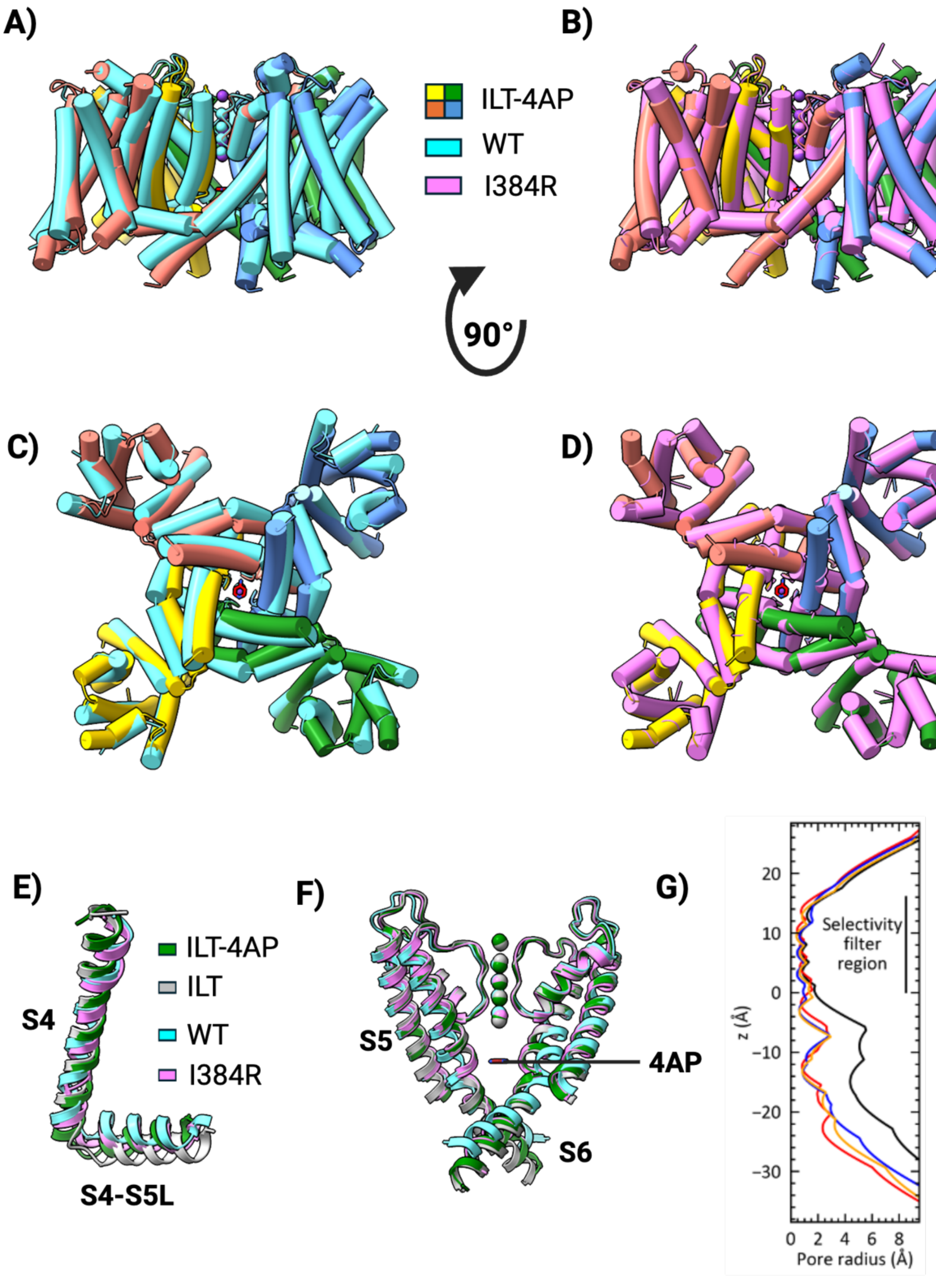


**Fig: S5. Conformational changes in Shaker ILT-4AP compared to WT and I384R.** A) & B) Membrane and intracellular view of Shaker ILT-4AP (salmon, cornflower blue, green and gold) compared with Shaker WT (teal). C) & D) Membrane and Intracellular view of Shaker ILT-4AP compared with Shaker I384R (pink). E) Comparison of S4 and S4-S5 linker among structures. F) Comparison of S5 and S6 helices among structures. G) Pore radius of ILT (red), ILT-4AP (Yellow), WT (Black) and I384R (Blue).

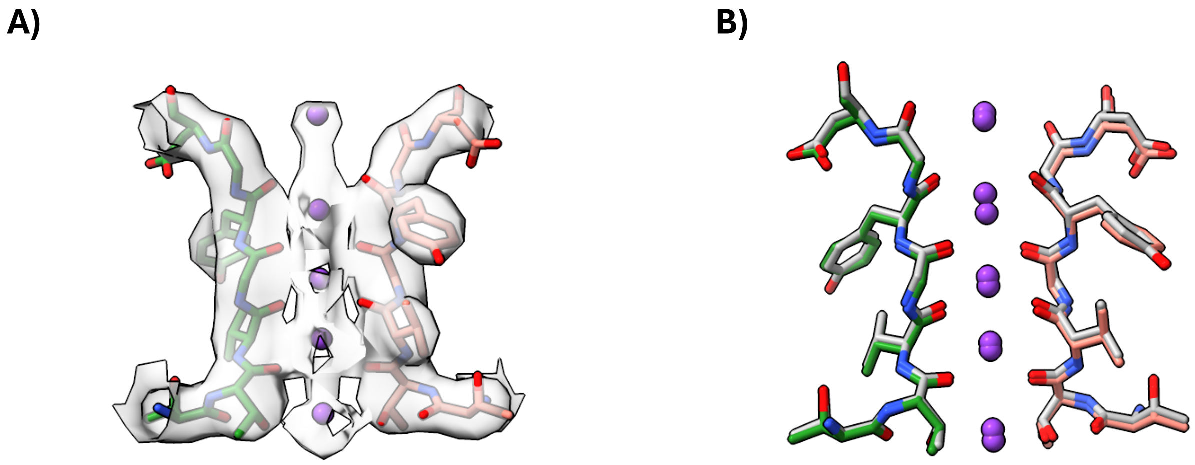

**Fig: S6. Selectivity filter comparison of Shaker ILT-4AP and ILT.** A) Shaker ILT-4AP model within unsharpened map showing selectivity filter region from T441 to D447 of opposite chains; $K^+$ map density is shown from sharpened map. B) Selectivity filter region from T441 to D447 of two chains from Shaker ILT-4AP (salmon & green) and Shaker ILT (gray); K+ ions colored in purple.

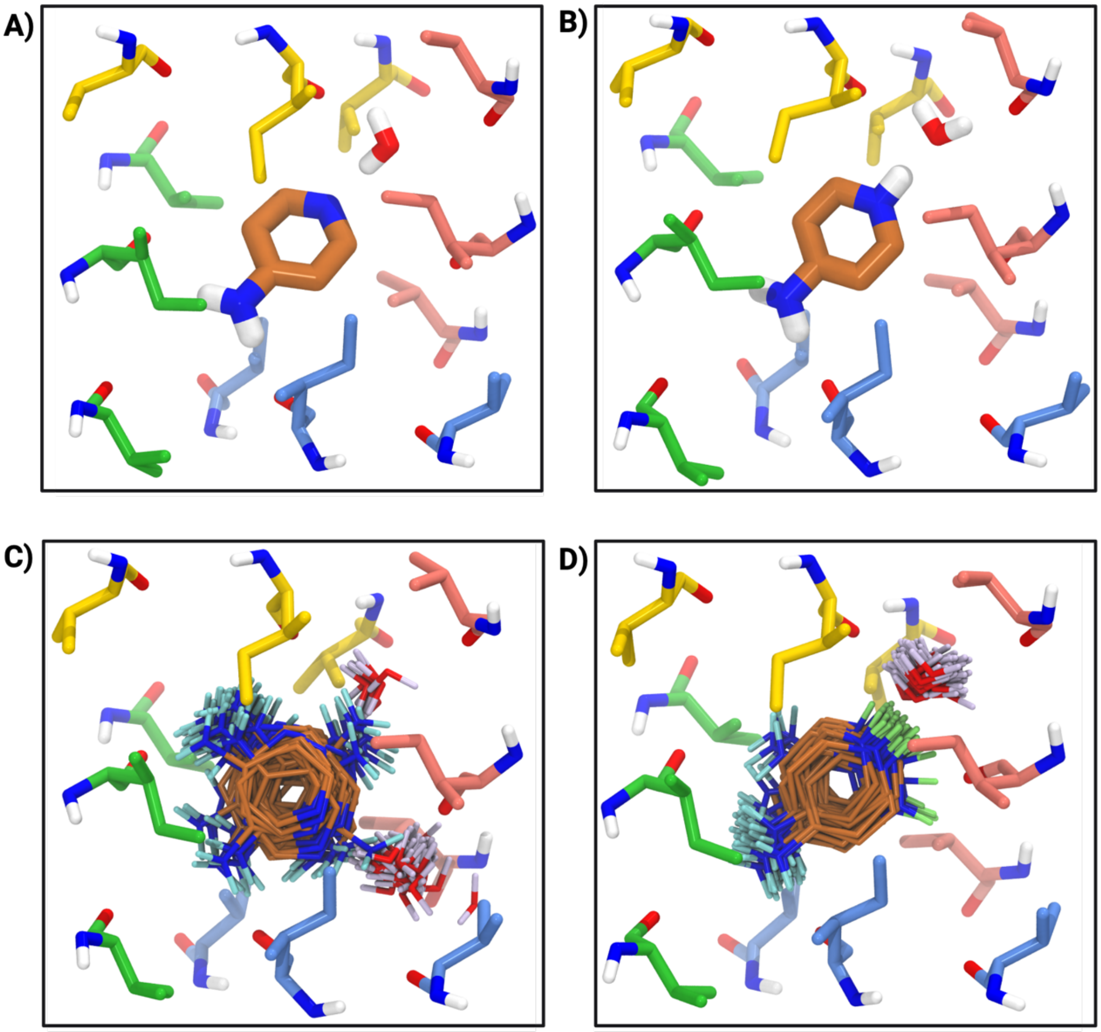


**Fig: S7. MD neutral (A, C) or protonated (B, D) 4AP bound to the closed channel.** Representative bound conformations (A, B) are shown for each protonation state. C, D shows an overlay of poses sampled by each state over 200 ns, hydrogens are colored for clarity. Four different colors (salmon, cornflower blue, green and gold) represent residues from different chains. 4AP ligand in brown color.

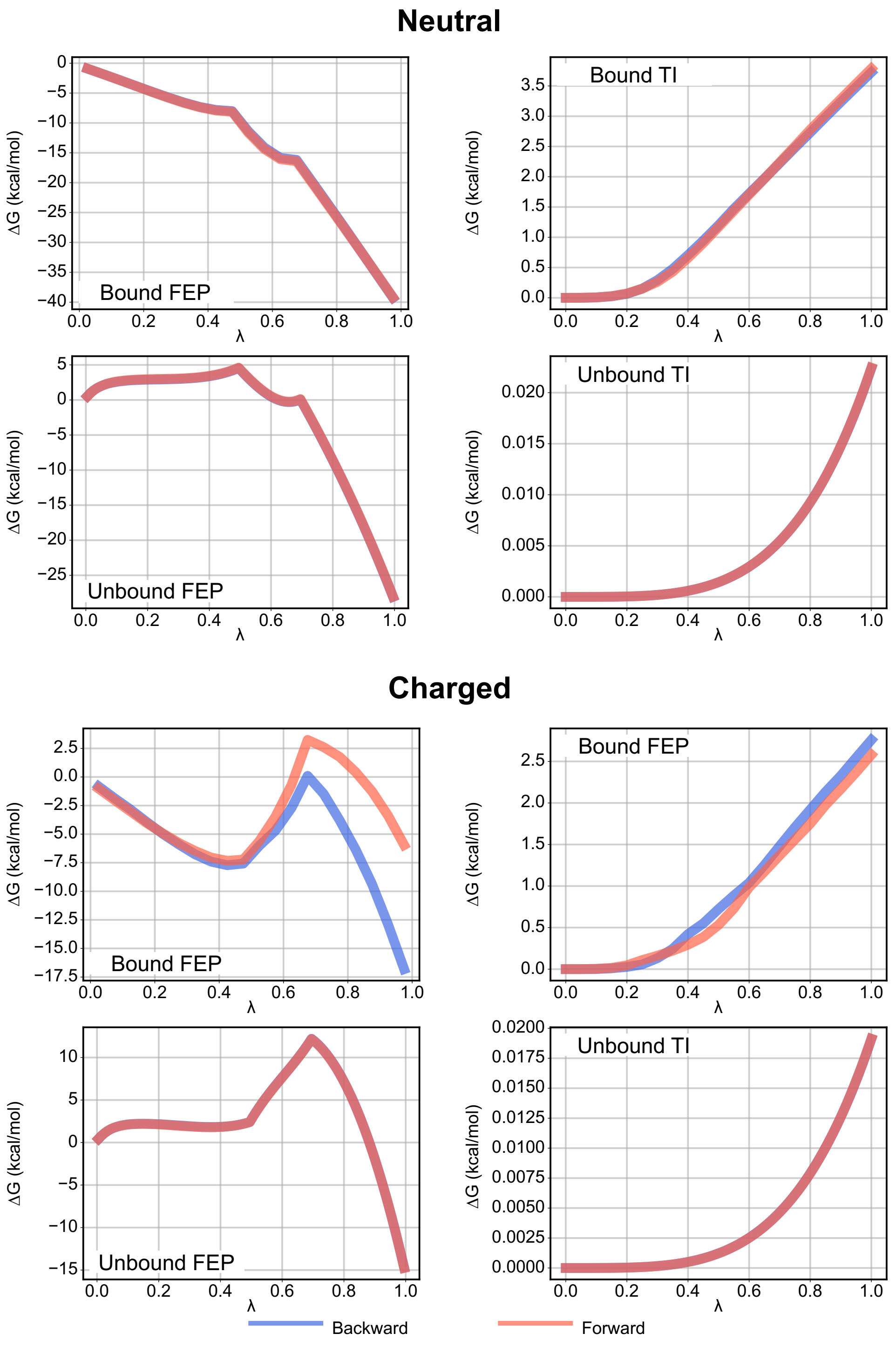


**Fig: S8. Change in free energy over coupling constant lambda**, which controls the progress between the decoupled (λ = 0) and coupled (λ = 1) states for the non-bonded forces during the free-energy perturbation (FEP) steps, or the restraints during the thermodynamic integration (TI) steps. Shown for the backward (blue) and forward (red) pathways. The large hysteresis in the Charged bound FEP is due to the loss of a binding water.

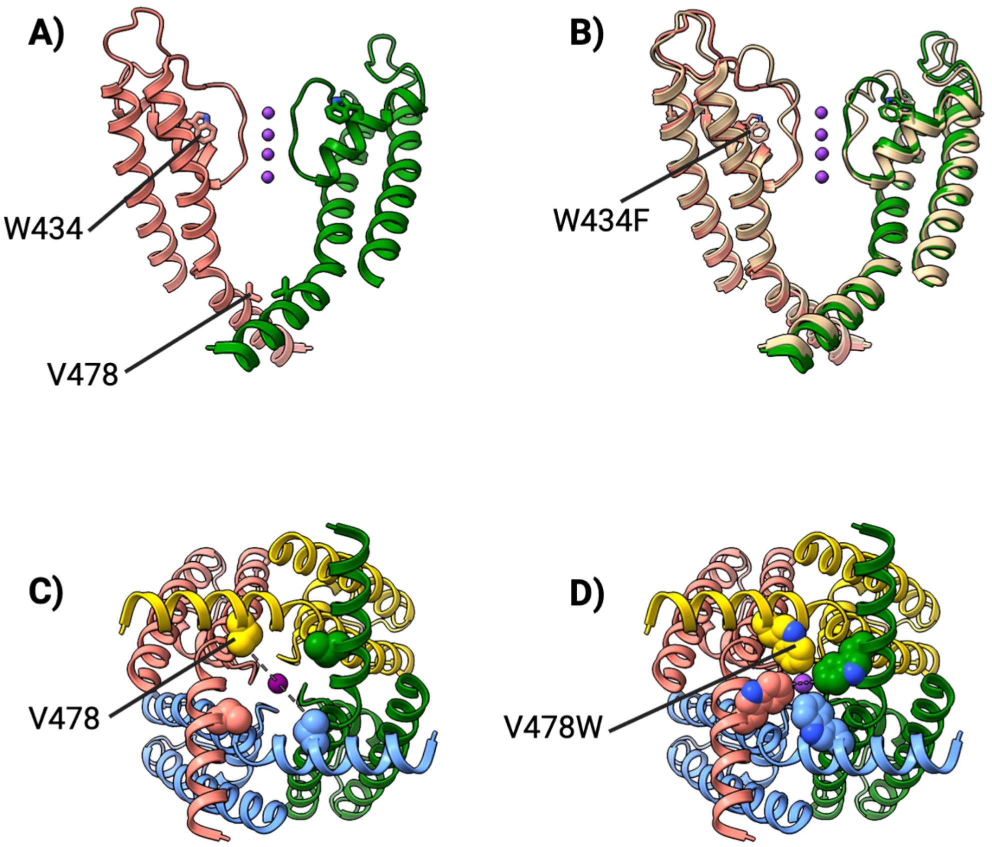


**Fig: S9. Effect of nonconductive mutant W434F and V478W. A)** Permeation pathway in the WT where positions of W434 and V478 are highlighted. **B)** Membrane view of permeation pathway showing 2 pore domains of WT (PDB ID 7SIP; salmon and green) and W434F (PDB ID 7SJ1; wheat) depicting dilation of selectivity filter. **C) and D)** Intracellular view of permeation pathway with **C)** position of V478 highlighted and **D)** mutated to W modified in chimeraX to show closed structure. Four different colors (salmon, cornflower blue, green and gold) represent different chains. K+ colored in purple.

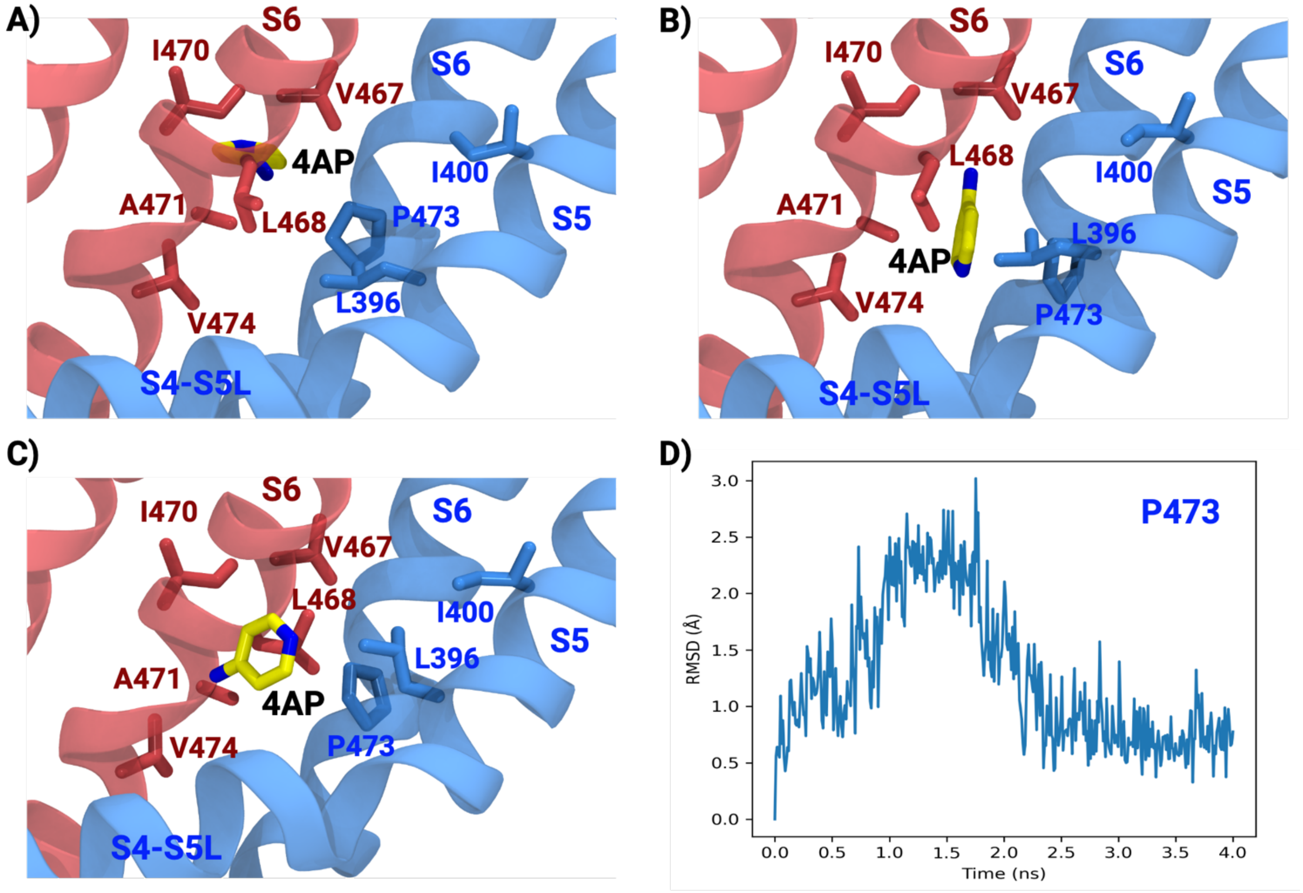


**Fig: S10. Molecular dynamics simulation of fenestration entry point.** A) B) and C) are three snapshots of MD simulations showing unbinding of 4AP. D) RMSD calculated for P473 of S6 helix.

A)

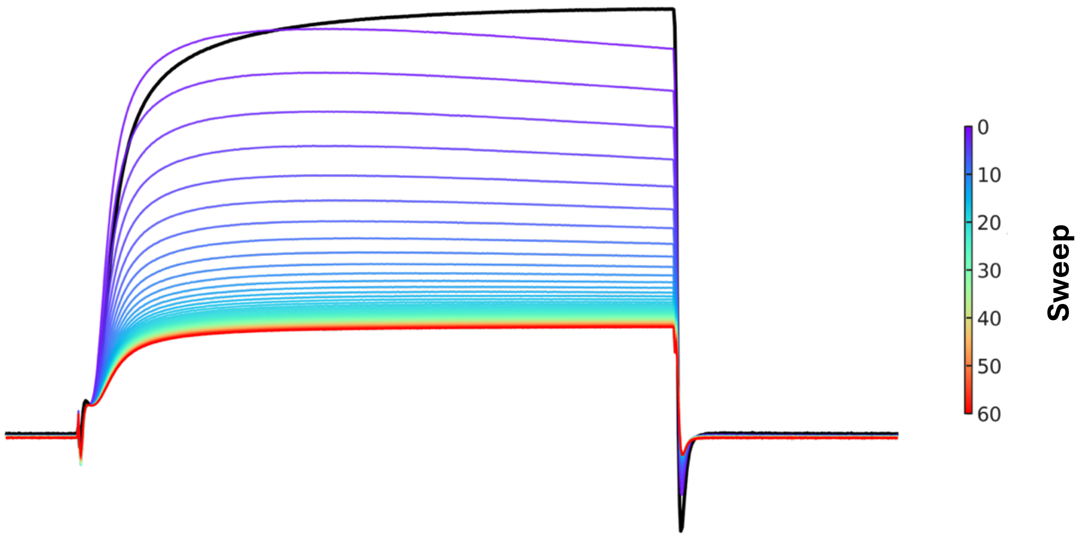


B)

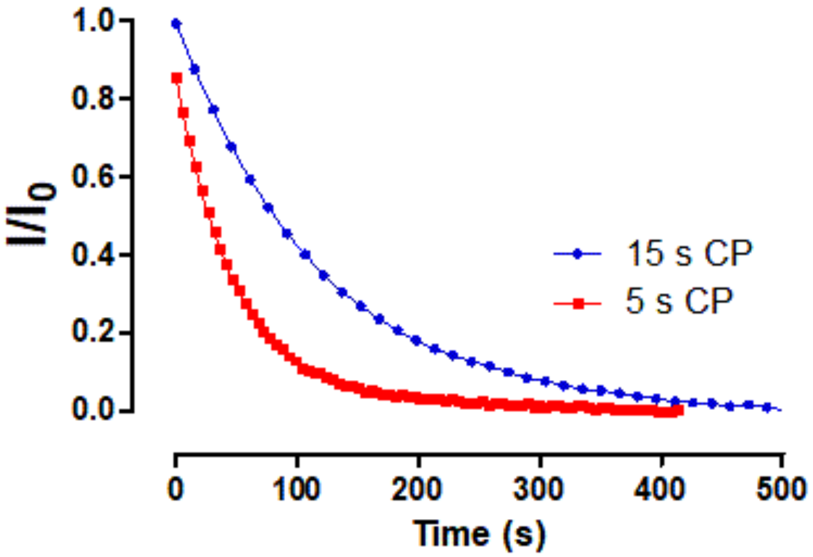


**Fig: S11. Determination of cycle period effect on inhibition kinetics. A)** Effect of repetitive activation with a cycle period (CP) of 5seconds, after 4AP application (400 µM) on the WT channel ionic currents. The black trace shows the currents before 4AP application. The discontinuous line indicates the time at which currents were measured to time course of 4AP inhibition. **B)** Inhibition time course after 4AP application when using a cycle period of 5 (red) or 15 (blue) seconds; continuous line corresponds to a single exponential decay fitting from where the time constants were extracted. Currents were normalized to the current before 4AP application minus the baseline after steady state.

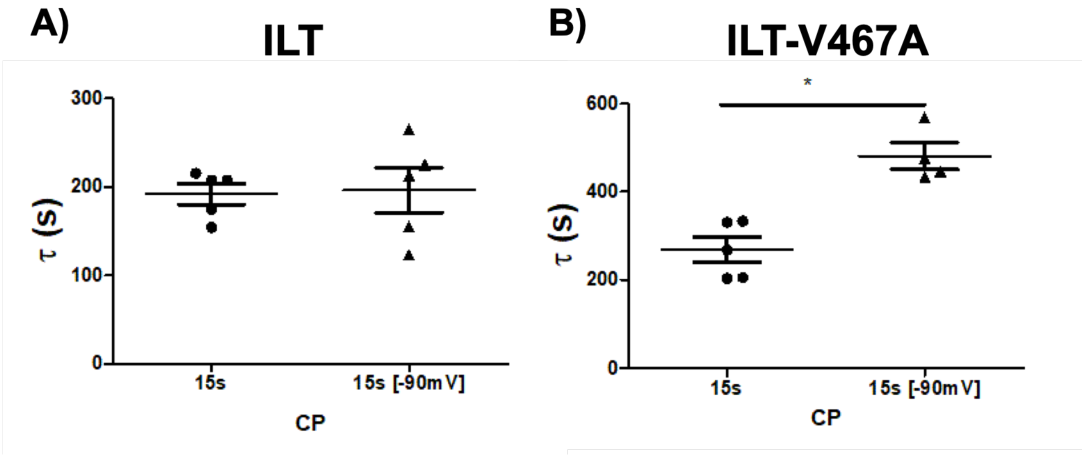


**Fig: S12.** Effect of holding voltage on the inhibition time constant of 4AP for (**A**) ILT (200 µM) and (**B**) ILT-V467A (1 mM) mutants. * $p<0.05$ Mann-Whitney U test.

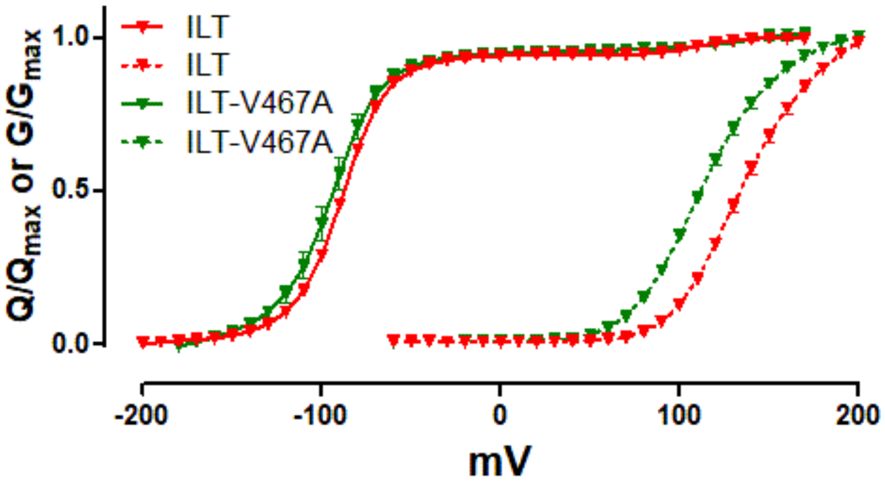


**Fig: S13. Voltage dependence of ILT-V467A mutant.** Q-V (solid lines) and G-V discontinuous lines) curves for the ILT (red) and ILT- V467A (green) mutants. Data Shown as mean ± SEM.

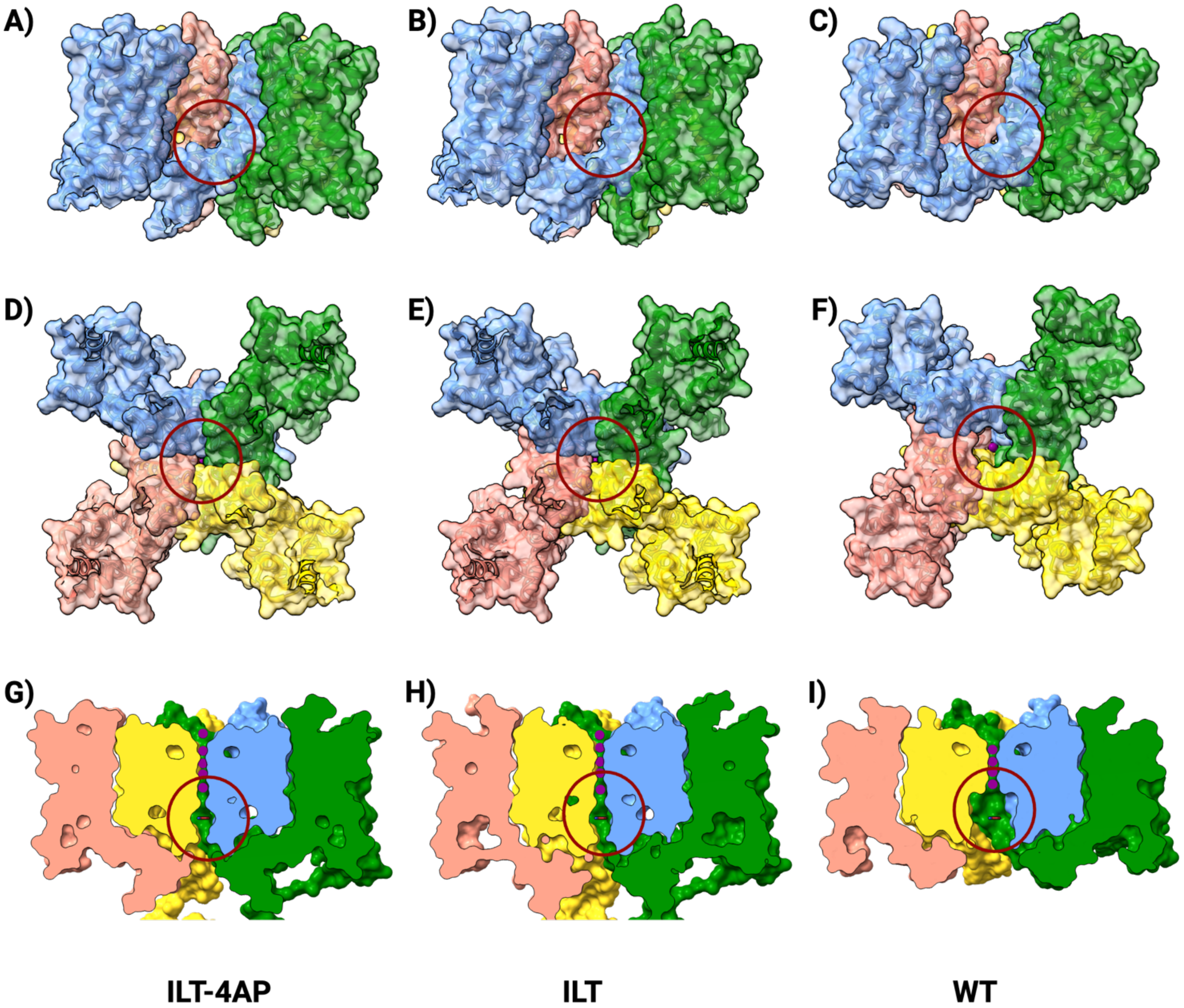


**Fig: S14. Possible entry points of 4AP into the pore in different structures.** Membrane view of **A)** ILT-4AP, **B)** ILT, and **C)** WT where red circle shows fenestration entry point. Intracellular view of **D)** ILT-4AP, **E)** ILT, and **F)** WT where red circle shows intracellular opening entry point. **G) H)** and **I)** Cross section to show 4AP binding pocket in ILT-4AP, ILT and WT respectively. Red circle shows 4AP binding pocket. Position of 4AP molecule in ILT and WT is overlapped from ILT-4AP. Chains A, B, C, D of all structures are colored in salmon, cornflower blue, green and gold.

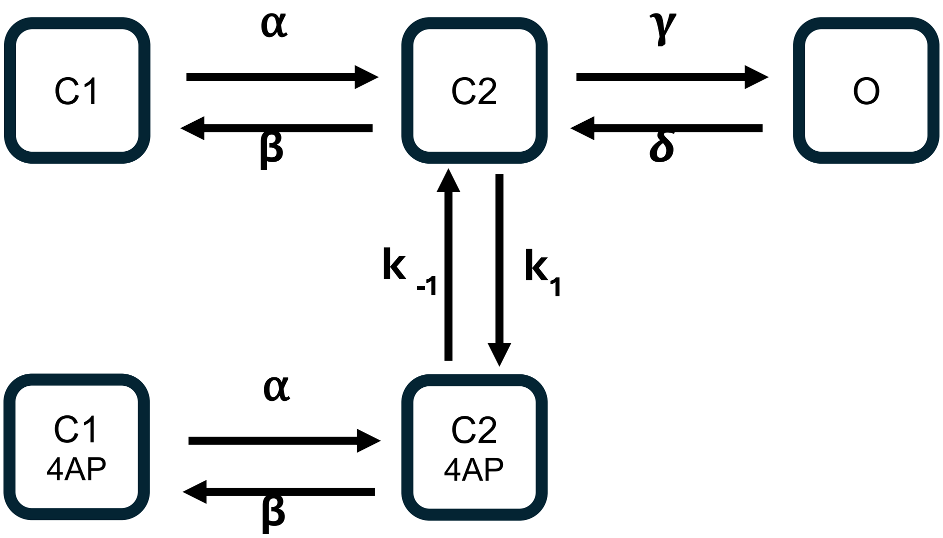


| Rate | Rate value ($s^{-1}$ or $(s*\mu M)^{-1}$) |
|---|---|
| α | $100\ e^{\frac{1.8*FV}{RT}}$ |
| β | $25\ e^{\frac{-1.8*FV}{RT}}$ |
| γ | $1000\ e^{\frac{0.2*FV}{RT}}$ |
| δ | $250\ e^{\frac{-0.2*FV}{RT}}$ |
| $k_1$ | $0.1 * [4 - AP]$ |
| $k_{-1}$ | $10$ |

**Fig: S15. Kinetic model details.** Diagram and parameter values for the 5-state 4AP bind kinetic model rates.
